# Unraveling Network Dynamics via RONI: Riemannian filtering Of Network Interactions

**DOI:** 10.1101/2025.03.26.645425

**Authors:** Yonatan Kleerekoper, Mohammad Kurtam, Yonatan Keselman, Shai Abramson, Dori Derdikman, Yitzhak Schiller, Simon Musall, Hadas Benisty

## Abstract

Functional connectivity (FC) is fundamentally non-stationary, undergoing continuous reconfigurations that track shifting behavioral and cognitive states. Despite the importance of these transitions, existing analytical frameworks struggle to reconcile the high-dimensional nature of these reconfigurations with the need for structured trajectories and mechanistic interpretability. Specifically, identifying the precise network components responsible for driving these dynamics remains a significant challenge. To bridge this gap, we introduce RONI (Riemannian filtering Of Network Interactions), a geometry-aware, unsupervised framework that treats time-varying FC as a signal evolving on the manifold of symmetric positive semidefinite (SPSD) matrices. By defining intrinsic filtering operators along the SPSD geodesics, RONI enables a principled multiresolution decomposition of FC dynamics directly on the manifold. This representation allows for the isolation of “dynamic drivers” - specific network elements that dominate coherent connectivity modes across distinct temporal scales. We demonstrate the versatility of RONI by applying it to a diverse array of large-scale neural recordings, including hippocampal electrophysiology data, cortex-wide and dendritic calcium imaging, and human EEG. Across these varied modalities and spatio-temporal scales, RONI identifies biologically meaningful sub-networks that shape the geometry of the connectivity trajectory, providing a unified, interpretable, and quantitative framework for studying the evolution of distributed neural interactions.

## I. INTRODUCTION

Understanding how brain connectivity reorganizes over time is a longstanding goal in systems and computational neuroscience [1–5]. Functional connectivity (FC), typically estimated through correlations between neural activity, is widely used to relate network interactions to behavior, state, and cognition [6–9].

FC is often analyzed edge by edge, treating correlations as a collection of pairwise values rather than as a structured matrix encoding population-level organization [10, 11]. Moreover, many traditional analyses assume FC is effectively stationary within a behavioral condition, averaging activity over long epochs and thereby neglecting its intrinsically dynamic nature [8, 10, 12, 13]. Such averaging collapses rich temporal structure into a single summary and can obscure transient, task-related, or learning-induced reconfigurations of network organization.

A common alternative, particularly in fMRI, is to model FC as a time-varying graph whose edges fluctuate over time [14–24]. While effective for capturing certain aspects of network dynamics, these approaches often emphasize individual edges or summary graph statistics, and may therefore miss coordinated, low-dimensional reconfigurations expressed in the joint structure of the full FC matrix [25].

A growing body of evidence suggests that FC dynamics form structured trajectories linked to changes in brain state, learning, and pathology [26–28]. Importantly, correlation matrices are symmetric positive semidefinite (SPSD) and thus do not form a Euclidean vector space: operations such as averaging, interpolation, and filtering must preserve both positivity and symmetry. Riemannian geometry provides principled tools for data-driven analysis of such data and for extracting low-dimensional trajectories. This technique has been used to study FC dynamics in calcium imaging [29, 30] and fMRI [31], but it does not provide a straightforward mechanistic interpretation.

Relatedly, supervised learning on Riemannian manifolds has been applied to EEG (e.g., brain–computer interface pipelines) [32–35] and to fMRI [36–38]. While these methods can improve predictive performance and offer some interpretability, their reliance on labels limits their ability to uncover intrinsic, data-driven mechanisms underlying FC reorganization. Interpretable and unsupervised Riemannian approaches to functional connectivity have also been proposed. Still, they are typically designed to compare connectivity across discrete states or conditions rather than to analyze continuous, multiscale sequences of FC matrices [39–41].

A notable step toward an interpretable and unsupervised framework for FC dynamics that respects SPSD geometry was introduced by [42]. Their approach is based on two operators defined on SPSD matrices: a similarity operator that preserves shared structure and a difference operator that captures deviations. However, while the similarity operator preserves the SPSD property, the difference operator does not. Because its output does not remain within the SPSD cone, the framework supports iterative smoothing. Still, it does not yield a complete multiresolution decomposition, thereby limiting its ability to resolve dynamics across temporal scales.

In this work, we introduce RONI (Riemannian filtering Of Network Interactions), a geometric, interpretable, and unsupervised framework for analyzing FC as a high-dimensional temporal signal evolving on the cone of SPSD matrices. Our central contributions are: (i) a novel high-pass operator that remains within the SPSD cone, (ii) a recursive filtering framework for extracting a multiresolution representation for FC dynamics, and (iii) an algorithm that attributes frequency-resolved connectivity changes to specific components of the underlying neural signals, which we term *dynamic drivers*. This construction yields a principled multiresolution decomposition that is both scalable and geometrically consistent. Crucially, by operating directly on SPSD matrices, RONI preserves the structure of FC throughout the analysis and links dominant frequency-specific modes back to the original neural components, providing interpretability that is often absent in unsupervised geometric embeddings.

We present a rigorous mathematical formulation of the framework together with an algorithm for detecting dynamic drivers. We apply RONI to large-scale neural recordings spanning multiple spatio-temporal scales, from cortical networks to dendritic dynamics, and multiple modalities, including calcium imaging, electrophysiology, and EEG. Across all modalities, RONI consistently recovers interpretable and biologically meaningful drivers of functional connectivity dynamics.

## II. METHODS

### A. Problem formulation

Let **x**[*t*] ∈ ℝ^*d*^ denote a multivariate time series of neural activity. We assume that the underlying FC of the network evolves slowly in time, so that pairwise interactions between the components of **x**[*t*] are approximately constant within a short window of *T*_*w*_ samples. We slide this window along the recording and compute the sample correlation in each window, yielding a sequence of SPSD matrices, 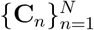, where *n* indexes window centers. Each **C**_*n*_ is SPSD, and the sequence {**C**_*n*_} therefore traces a trajectory on the SPSD manifold. This trajectory describes the FC temporal dynamics of the observed network. This setup gives rise to two central questions:

1. How can we characterize the temporal dynamics of the FC sequence, 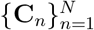 while respecting the manifold geometry?
2. Which components of **x**[*t*] drive these FC dynamics?

A commonly used approach for the first question is to apply nonlinear manifold learning to obtain a low-dimensional coordinate system that preserves the intrinsic geometry of correlation matrices [35, 36, 43]. We use diffusion maps [44] with affinities defined by Riemannian distances on the SPSD manifold. Specifically, we compute pairwise distances *d*(**C**_*n*_, **C**_*m*_) using the geodesic-distance extension for SPSD matrices [45] (see Appendix V A for more details) and define the diffusion kernel

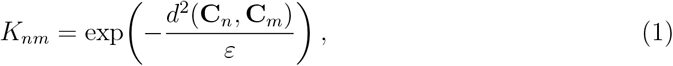

where *ε* > 0 is the kernel bandwidth. Let **D** be the diagonal matrix with 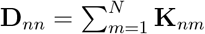 and define the row-stochastic diffusion operator **P** = **D**^−1^**K**. Denoting by 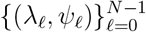 the eigenpairs of **P** (ordered as 1 = *λ*_0_ > *λ*_1_ ≥ · · · ≥ 0), the diffusion-maps embedding is

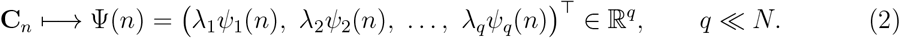

This approach has been used to obtain low-dimensional representations of time-varying FC and relate them to behavior or learning [12, 29, 30].

The second question is more challenging: geometry-aware methods (including diffusion maps) typically provide no explicit inverse mapping from the low-dimensional coordinates Ψ(*n*) back to specific entries of **C**_*n*_, and thus do not directly identify which components of **x**[*t*] drive the observed FC changes. Our approach, RONI, addresses both questions simultaneously, providing a multi-resolution decomposition of 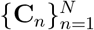 and detecting components of **x**[*t*] that consistently dominate in informative frequency bands, which we term *dynamic drivers*.

### B. Similarity and Difference operators on the SPSD manifold

The core building blocks of our method, RONI, are two geometric operators that act on pairs of SPSD matrices: a *similarity* operator, which emphasizes shared structure, and a *difference* operator, which accentuates changes. Both are defined via the affine-invariant Riemannian geodesic connecting two matrices.

Let **C**_1_ and **C**_2_ be SPD. The geodesic *γ*(*p*) satisfying *γ*(0) = **C**_1_ and *γ*(1) = **C**_2_ is

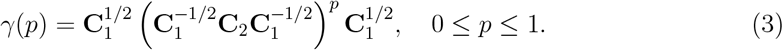

Bonnabel et al. [45] extended this construction to SPSD matrices; we denote the resulting SPSD geodesic by 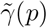 (see Appendix V A). In particular, they prove the following:

#### Proposition 1.

For any pair of SPSD matrices **C**_1_ and **C**_2_, the geodesic 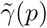 is well-defined for all *p* ∈ R. Moreover, the SPSD manifold is geodesically complete [45].

Following Shnitzer et al., the similarity operator is defined as the geodesic midpoint:

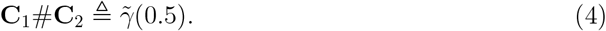

This midpoint operation emphasizes the structure shared between **C**_1_ and **C**_2_.

We next introduce a *difference operator* that extrapolates along the same geodesic beyond **C**_2_:

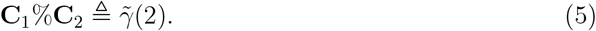

Proposition 1 guarantees that for any given SPSD inputs, the outcome is also SPSD; therefore, the operator preserves the manifold structure. Fig. 1A presents a geometric illustration of the difference operator as it extends the geodesic so that **C**_2_ becomes the midpoint between **C**_1_#**C**_2_ and **C**_1_%**C**_2_. The difference operator is not symmetric: in general, **C**_1_%**C**_2_≠ **C**_2_%**C**_1_. The two operators # and % are mutually inverse (Proposition 2, Appendix V D) and form the basis for our geometric filtering scheme. In the next section, we show that the difference operator selectively amplifies eigenmodes that differ between **C**_1_ and **C**_2_, thereby emphasizing changes in FC structure.

**FIG. 1.**
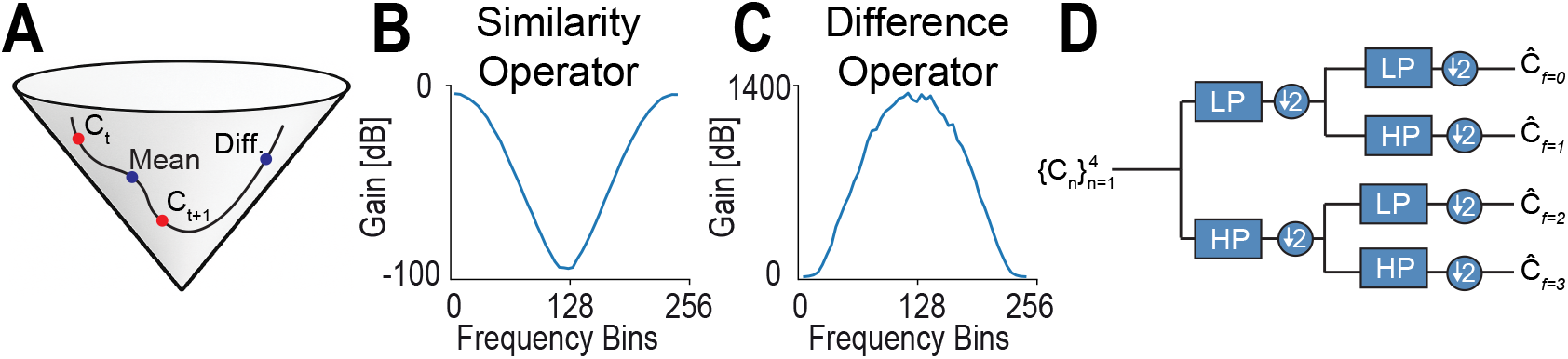
(A) A schematic illustration of the geodesic connecting two matrices where the similarity operator is positioned at the midpoint, while the difference operator extends the geodesic such that **C**_2_ becomes the midpoint between **C**_1_%**C**_2_ and **C**_1_. (B) Frequency response of the low-pass filter. (C) Frequency response of the high-pass filter. (D) An illustration of the proposed filtering framework: recursive application of low-pass and high-pass filters, followed by down-sampling by a factor of two.

### C. Spectral analysis

This section characterizes how the similarity (#) and difference (%) operators transform the *spectral structure* (eigenvalues and eigenvectors) of their inputs. When **C**_1_ and **C**_2_ are FC estimates from adjacent time windows, we expect many large-scale network modes to remain approximately stable, while changes in FC may appear as changes in the *strength* of particular modes. We formalize this intuition by analyzing the action of # and % on shared (or approximately shared) eigenmodes.

We begin with an idealized case in which **C**_1_ and **C**_2_ share an eigenvector *ψ* but assign it different eigenvalues.

#### Theorem 1

*Let ψ be an eigenvector of both* **C**_1_ *and* **C**_2_ *with eigenvalues λ*_1_ *and λ*_2_, *respectively:* **C**_1_*ψ* = *λ*_1_*ψ*, **C**_2_*ψ* = *λ*_2_*ψ. Then ψ is also an eigenvector of the similarity operator* **C**_1_ #**C**_2_, *with eigenvalue:* 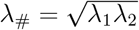.

Theorem 1 (follows from prior work by [42]) shows that the similarity operator preserves shared eigenmodes while combining their strengths via a geometric mean, consistent with its interpretation as a geometry-aware “averaging” operation.

We next state the corresponding result for our difference operator, which is central to isolating changes in FC.

#### Theorem 2

*Under the same assumptions of Theorem 1 and with λ*_1_ > 0, *ψ is also an eigenvector of* **C**_1_%**C**_2_, *with eigenvalue:* 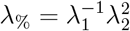.

*Proof*. See Appendix V B.

Theorem 2 reveals how % emphasizes changes in mode strength: the mapping 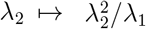 amplifies increases in *λ*_2_ relative to *λ*_1_ and suppresses decreases. In other words, if a coordinated network pattern *ψ* becomes more prominent from one window to the next (*λ*_2_ > *λ*_1_), then applying **C**_1_%**C**_2_ accentuates that mode. This property underlies our use of % to isolate temporally evolving connectivity features and, ultimately, to attribute those changes to specific neural components - dynamic drivers.

Exact eigenvector sharing is a strong assumption. In practice, # and % are applied to matrices from adjacent time windows where eigenspaces might be only approximately aligned. Appendix V B provides a perturbation analysis showing that the above spectral behavior is stable under small deviations, supporting the robustness of these operators for slowly varying FC.

### D. Similarity and difference operators as low- and high-pass filters

We now leverage the spectral properties in Sec. II C to define intrinsic, Riemannian analogs of low-pass and high-pass filtering for an SPSD matrix sequence. For consecutive FC matrices, we define the low-pass and high-pass outputs as

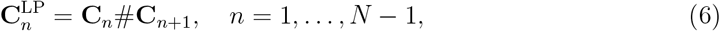

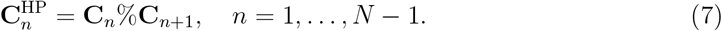

The similarity operator # computes the geodesic midpoint, so 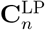provides a geometrical smoothing of FC across adjacent windows. In contrast, the difference operator % accentuates deviations between **C**_*n*_ and **C**_*n*+1_, acting as a high-pass operation on connectivity dynamics.

Classical filters for 1-dimensional signals are characterized by their frequency response, defined as the relative output amplitude obtained when the input is a sinusoid at a single frequency. We adopt the same operational definition here. We probe the SPSD filters with a controlled input sequence in which only a small subnetwork exhibits sinusoidal connectivity dynamics at frequency *f*, while all remaining entries are unstructured noise. Specifically, we generate a sequence 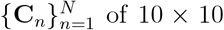 SPSD matrices such that correlations within components 1–3 oscillate sinusoidally,

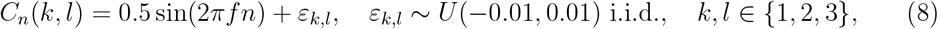

and all other off-diagonal entries are noise, with unit diagonal:

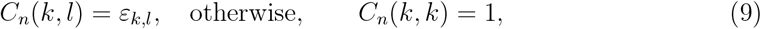

where *U* (*a, b*) denotes the uniform distribution on [*a, b*].

To quantify an “amplitude” for the filtered matrix sequence, we exploit that the structured oscillation is low-dimensional and is expressed primarily through the dominant spectral mode of **C**_*n*_. We therefore compute, at each time index *n*, the fraction of total variance explained by the leading eigenvalue,

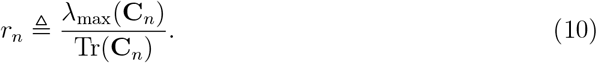

Large *r*_*n*_ indicates that a single coherent connectivity mode dominates the matrix, consistent with a structured subnetwork contribution, whereas smaller *r*_*n*_ indicates a more spectrally diffuse (noise-like) matrix.

We summarize the sequence by the average complement of this dominance ratio,

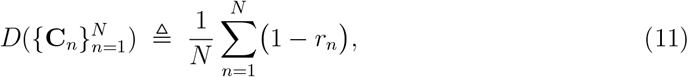

and refer to *D*(·) as a *spectral dispersion index*. Finally, given an input sequence 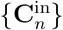 and its filtered output 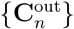, we define the frequency response at frequency *f* as the ratio of dispersion indices (in dB),

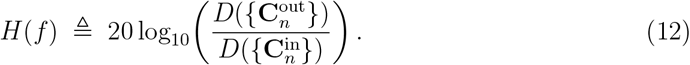

With this construction, *H*(*f*) reports how strongly the filter attenuates (or preserves) structured sinusoidal connectivity dynamics at frequency *f* . Fig 1B and C show the frequency responses of the resulting low- and high-pass filters, computed numerically from *N* = 256 matrices generated according to Eq. (8). As expected, the low-pass filter suppresses higher frequencies, whereas the high-pass filter suppresses lower frequencies.

### E. RONI: Riemannian filtering Of Network Interactions

In this section we introduce our proposed framework, RONI, that captures time-varying functional connectivity across multiple temporal scales and identifies network components that consistently drive structured changes in connectivity. Starting from an SPSD FC sequence 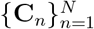, RONI performs a geometry-preserving multi-resolution decomposition on the SPSD cone and then extracts *dynamic drivers* by aggregating evidence across frequency-resolved connectivity modes.

#### Riemannian multiresolution filtering on the SPSD manifold

Building on the Riemannian low-pass and high-pass operators in Eqs. (6)–(7), we construct a multiresolution decomposition analogous to a wavelet-packet transform for scalar signals, but computed intrinsically on the SPSD manifold. Given an input sequence of *N* SPSD correlation matrices, at the first scale, we apply the low-pass and high-pass filters to adjacent pairs and then downsample by a factor of two. This yields two SPSD sequences of length *N/*2: the low-pass branch captures slowly varying FC structure shared across adjacent windows, whereas the high-pass branch captures faster FC changes emphasized by the difference operator. We then apply the same filtering-and-downsampling procedure again to each branch. This yields four SPSD sequences, each of length *N/*4, corresponding to a finer partition of four frequency bands and a reduction in temporal resolution by an additional factor of two (see Fig. 1D). For *N* a power of two, we recursively iterate *log*_2_*N* steps of filtering and downsampling, till each final node contains a single SPSD matrix. We denote these terminal coefficients by 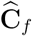, where the index *f* labels the frequency bin associated with that terminal node. At this maximal depth, the representation attains the finest frequency resolution, while temporal evolution is no longer represented within a bin.

#### Dynamic drivers detection from frequency-resolved connectivity modes

Each coefficient 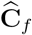 summarizes FC structure concentrated within frequency bin *f* . For each bin we compute the dominant eigenpair

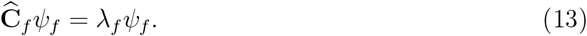

and interpret *ψ*_*f*_ as the leading frequency-specific connectivity mode. In particular, |*ψ*_*f*_ [*i*]| reflects the contribution of component *i* to that dominant mode.

Not all frequency bins are informative: some coefficients are dominated by diffuse, noise-like structure. To quantify whether *ψ*_*f*_ is concentrated on a small subset of components—consistent with a structured subnetwork—we compute the normalized entropy of its magnitude profile:

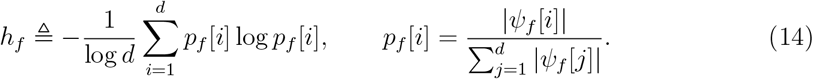

The normalization by log *d* ensures *h*_*f*_ ∈ [0, 1]. Low entropy indicates that the dominant mode is concentrated on a few components, whereas high entropy indicates a more distributed mode.

We retain a set of informative frequency bins using two criteria: (i) **concentration**, *h*_*f*_ ≤ *h*_cutoff_, and (ii) a **frequency-range constraint**, *f < f*_cutoff_, which biases the analysis toward structured, slower fluctuations and away from the highest-frequency bins that are more likely to reflect noise. Denoting the retained set by

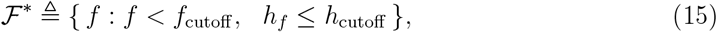

We next convert each retained eigenvector into a binary membership pattern over components. Specifically, for each *f* ∈ ℱ^∗^, we apply *k*-means clustering (*k* = 2) to the values 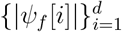, partitioning components into high- and low-amplitude groups. This yields an indicator **1**_*f*_ [*i*] ∈ {0, 1}, where **1**_*f*_ [*i*] = 1 if component *i* belongs to the high-amplitude cluster for bin *f* .

Finally, we aggregate evidence across frequencies by defining a voting score for each component,

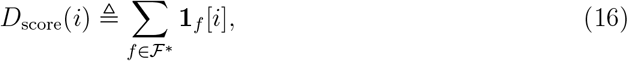

and identify *dynamic drivers* as those components whose score exceeds a threshold *M* :

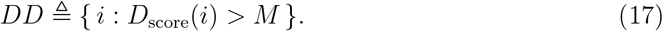

Dynamic drivers are thus components that repeatedly dominate leading connectivity modes across multiple temporal scales. The full procedure is summarized in Algorithm 1 (Fig. 2).

**FIG. 2.**
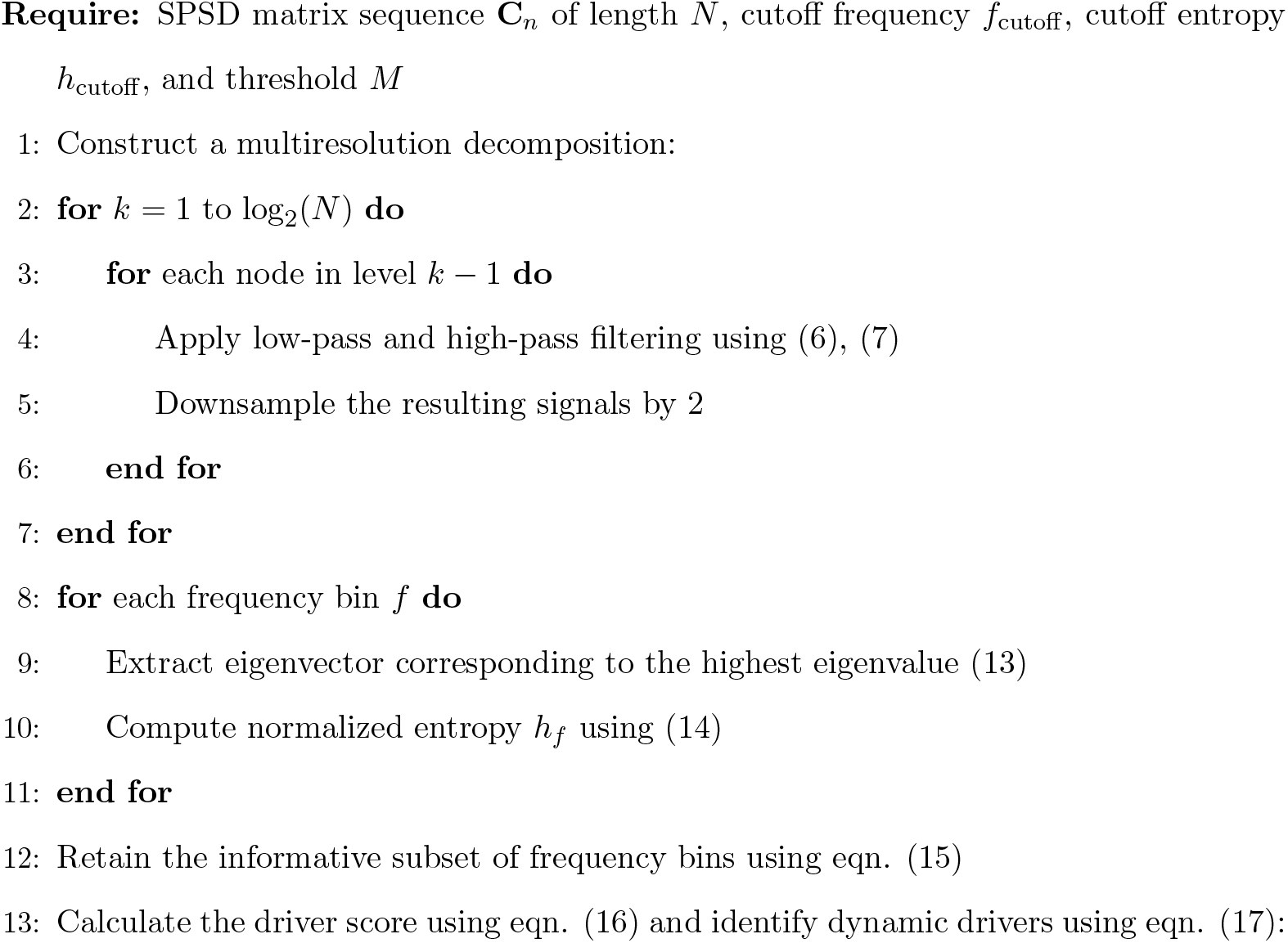
**Algorithm 1**: Dynamic drivers detection using RONI: Riemannian filtering Of Network Interactions.

We illustrate RONI on a controlled toy example that contains two temporally distinct subnetworks and background noise. Specifically, we generate a sequence of *N* = 256, 20 *×* 20 SPSD matrices 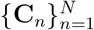 in which correlations among components 1–3 oscillate at a low frequency *f*_1_ = 4*/*256 *cycles/sample*, correlations among components 7–9 oscillate at a higher frequency *f*_2_ = 66*/*256 *cycles/sample*, and all other off-diagonal entries are unstructured noise (see Fig. 3A):

**FIG. 3.**
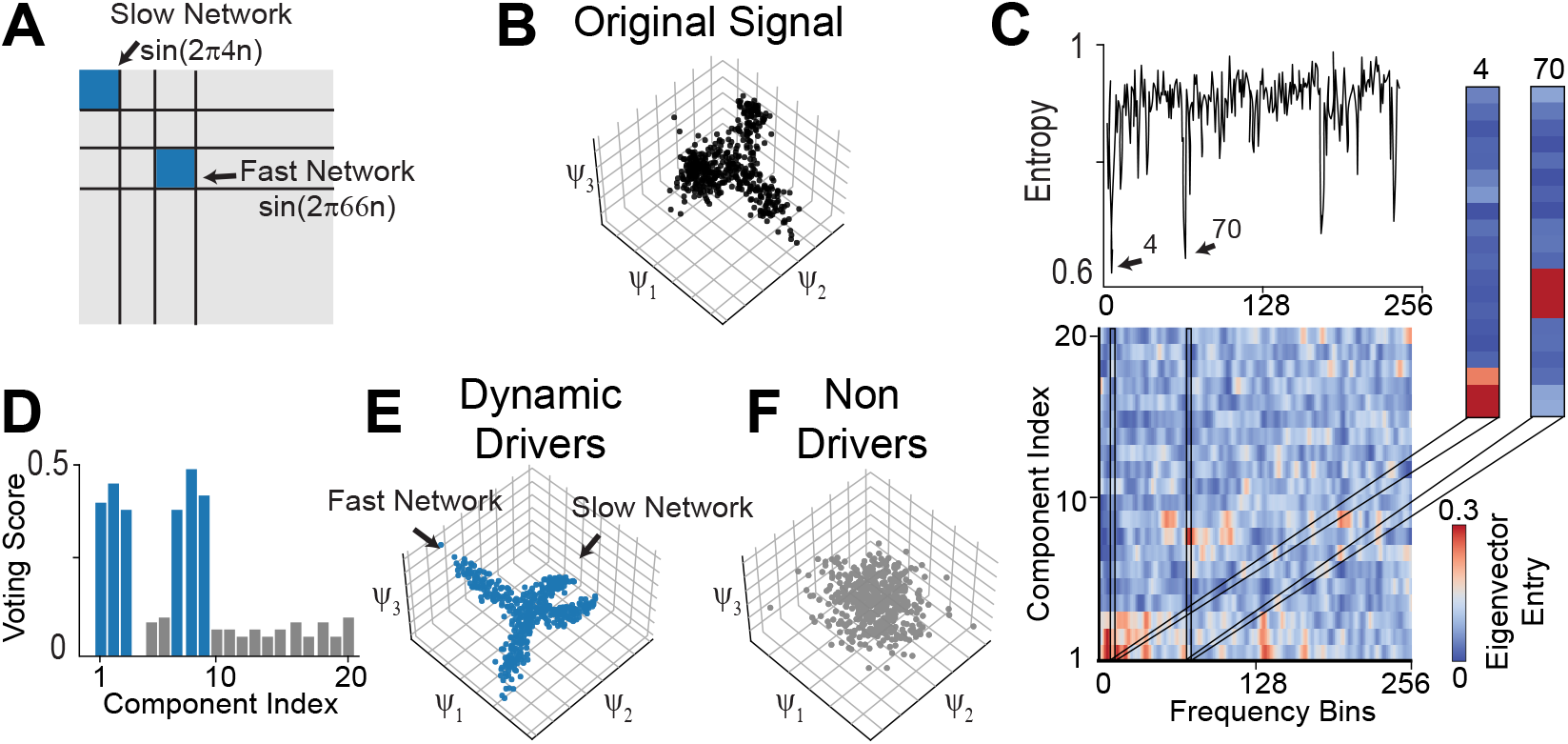
Toy example: a simulated sequence of SPSD matrices. (A) Schematic representation of the toy example entries, where entries 1–3 exhibit slow dynamics and entries 7–9 exhibit fast dynamics; gray areas indicate unstructured noise. (B) Diffusion embedding of the sequence, computed using all components. (C) (Top) Entropy computed across frequency bins. (Bottom) Entry values of the leading eigenvector across frequency bins. (Right) Leading eigenvectors associated with frequency bins 4 and 70, demonstrating that dynamic components are preferentially highlighted. (D) Voting scores across components, blue - true dynamic drivers, gray - non-drivers. (E,F) Diffusion embedding of the sequence of matrices considering: E. dynamics drivers, F. non-drivers only.

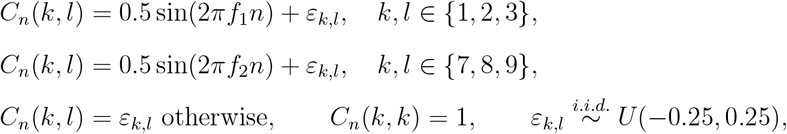

Fig. 3B shows the diffusion embedding of this sequence as a noisy cloud where the expected structure of two distinct domains corresponding to the two halves of the sequence is barely recognizable. Applying Algorithm 1 yields low-entropy leading eigenvectors in the frequency bins 4 and ∼ 70 respectively (Fig. 3C, top). The eigenvectors corresponding to these frequency bins point to components 1–3 and 7–9, respectively. Aggregating across informative bins via the voting score in Eq. (16) assigns the highest scores to components 1–3 and 7–9 (Fig. 3D), correctly identifying them as dynamic drivers.

To test whether the identified drivers preserve the intrinsic geometry of the FC dynamics, we compare the diffusion-map embeddings computed from the whole matrix sequence (Fig. 3B) to (i) a driver-restricted sequence obtained by retaining only rows and columns corresponding to detected drivers, and (ii) the complementary sequence containing only non-drivers. The driver-restricted sequence recovers a clean, low-dimensional manifold consistent with the generative structure (Fig. 3E), whereas the non-driver sequence yields an unstructured embedding (Fig. 3F). Together, these results demonstrate that RONI isolates components carrying structured multi-scale connectivity dynamics while suppressing unstructured noise.

## III. RESULTS

RONI provides an interpretable view of how functional connectivity (FC) reorganizes over time and which components predominantly shape these dynamics. We demonstrate the framework across multiple recording modalities and spatial scales, from large-scale multiarea electrophysiology and cortex-wide imaging to dendritic calcium signals and human EEG, highlighting both generality and interpretability.

### A. Hippocampal FC encodes spatial position through dynamic drivers

We first asked whether short-timescale FC dynamics within hippocampal–entorhinal circuits encode spatial position during navigation. We analyzed chronic Neuropixels recordings from freely moving mice (*n* = 4) exploring multiple arena geometries (38 arenas; 20 min per arena), targeting subiculum, medial entorhinal cortex (MEC), and temporal association cortex (TEA) (Fig. 4A,B; Appendix V F). We estimated firing rates in 0.25 s bins and computed short-term correlations using a sliding window (4 s length; 2 s hop), yielding a sequence of SPSD correlation matrices 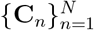 over each exploration session. Examples for firing rate traces along with the x-y position as a function of time, and as trajectories in three selective arenas are presented in Fig. 4A and B, respectively.

**FIG. 4.**
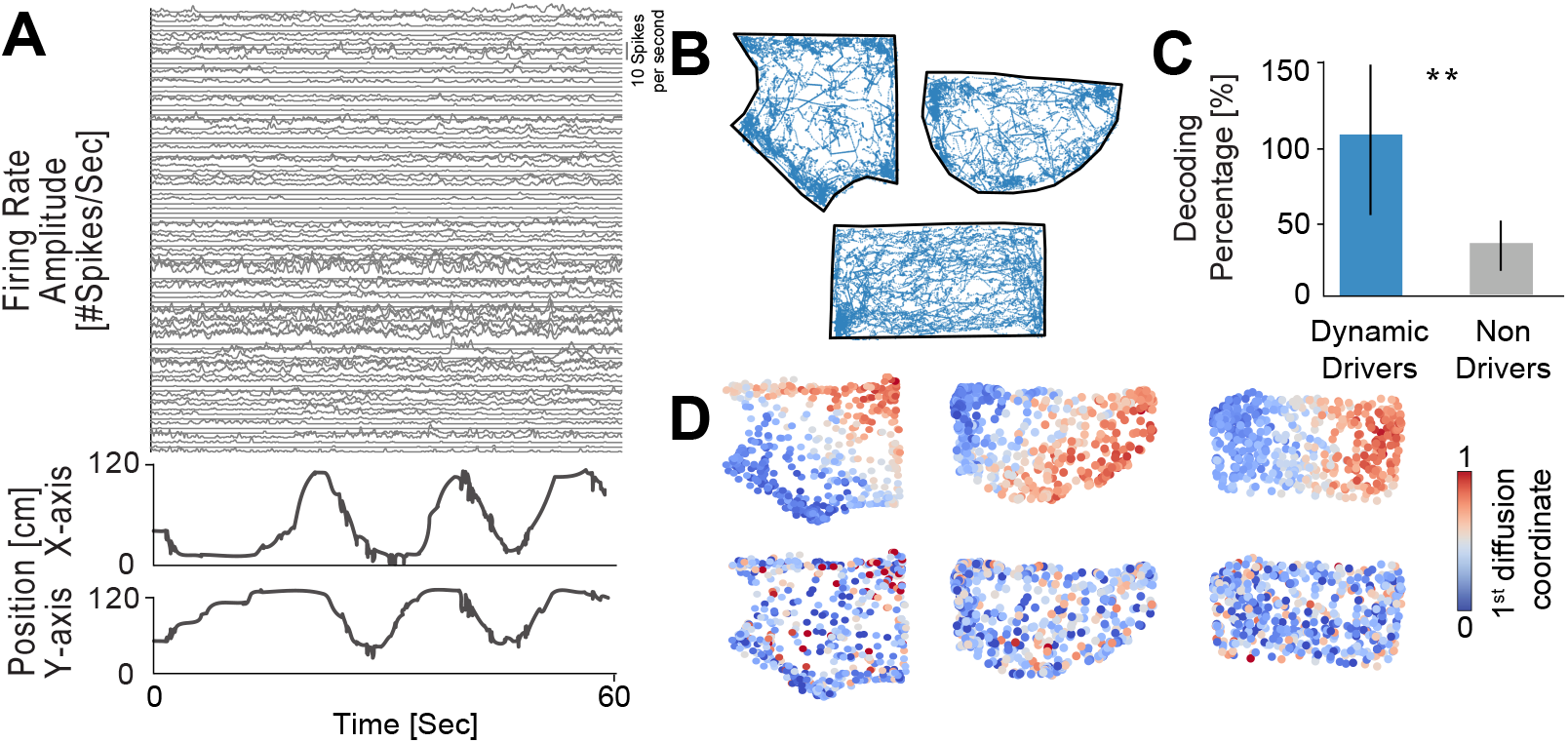
Spatially informative hippocampal dynamics. (A) Top: Example firing-rate traces of neurons during a 60s segment extracted from a 20 min recording session. Bottom: Corresponding mouse position trace during the same 60s interval. (B) Three example arena outlines (black) with the full mouse trajectories overlaid in blue. (C) Decoding power (percentage) calculated as the accuracy (*R*^2^) for predicting location from correlation dynamics using driving cells (blue) and non-driving cells (gray), normalized by the *R*^2^ achieved by the entire network (mean *±* SEM, *n* = 38 arenas). (D) Mouse (*x, y*) position colored by the first diffusion coordinate for drivers (top) and non-drivers (bottom) in three example arenas, illustrating that driver-restricted embeddings form smooth spatial gradients, whereas non-drivers lack structure.

For a detailed description of the recording setup and pre-processing, see Appendix V F. We computed short-term correlations using a sliding window (4-sec length; 2-sec hop), yielding a sequence of SPSD correlation matrices over each exploration session.

To quantify whether FC dynamics encode position, we embedded the FC trajectory using diffusion maps (see Methods, eqs. (1)–(2)) and trained a linear decoder to predict the animal’s (*x, y*) location from the 10 leading embedding coordinates. Decoding was robust across arenas (*R*^2^ = 0.35 ± 0.04, mean ± SEM; 10-fold cross-validation), indicating that the FC trajectory contains spatial information.

We then asked which neurons drive these informative FC dynamics. We applied RONI to each session and identified *dynamic drivers*. We retained frequency bins whose leading eigenvectors had the lowest entropy (with *h*_cutoff_ set to the median). We labeled neurons as drivers if their scores exceeded the population mean, with a cap of 50% drivers per session. We repeated the same embedding-and-decoding pipeline using only the driver subnetwork or only the complementary non-driver subnetwork. Fig. 4C presents the *R*^2^ obtained by each sub-population divided by the one obtained by the entire population (in percentage). Decoders built from driver-based FC dynamics significantly outperformed those built from non-drivers (*p* = 3.8 *×* 10^−6^), and achieved performance comparable to (and sometimes exceeding) the full population.

This difference was also apparent qualitatively: when coloring position by the first diffusion coordinate, driver-restricted embeddings formed smooth, spatially ordered gradients across arenas, whereas non-driver embeddings lacked organized structure (Fig. 4D). Together, these results show that navigation-related information in short-term FC dynamics is concentrated in a subpopulation of dynamic drivers that shapes the geometry of the FC trajectory.

### B. Learning-related cortical reorganization is dominated by a reproducible set of driver regions

Next, we used RONI to study learning-related reorganization of cortex-wide FC. We analyzed widefield calcium imaging data from mice performing an auditory decision-making task [46] (see Appendix V G for a detailed description of the experimental setup). We included four animals, each contributing ∼1,400 successful trials over ∼30 sessions, restricting analysis to successful detection trials to reduce behavioral confounds. Activity was parcellated into 19 cortical areas (one hemisphere; Allen CCF), and FC sequence was computed using a sliding window of 20 trials.

We embedded each animal’s FC trajectory using diffusion maps (Fig. 5A). We trained a linear decoder to predict session index from the embedded coordinates, treating session index as a proxy for training progression. We then applied RONI within each animal and asked whether driver-restricted FC trajectories better captured learning progression than the complementary non-driver set. Across animals, driver-based FC dynamics yielded higher decoding accuracy than the full network, whereas non-drivers consistently performed worse (Fig. 5B).

**FIG. 5.**
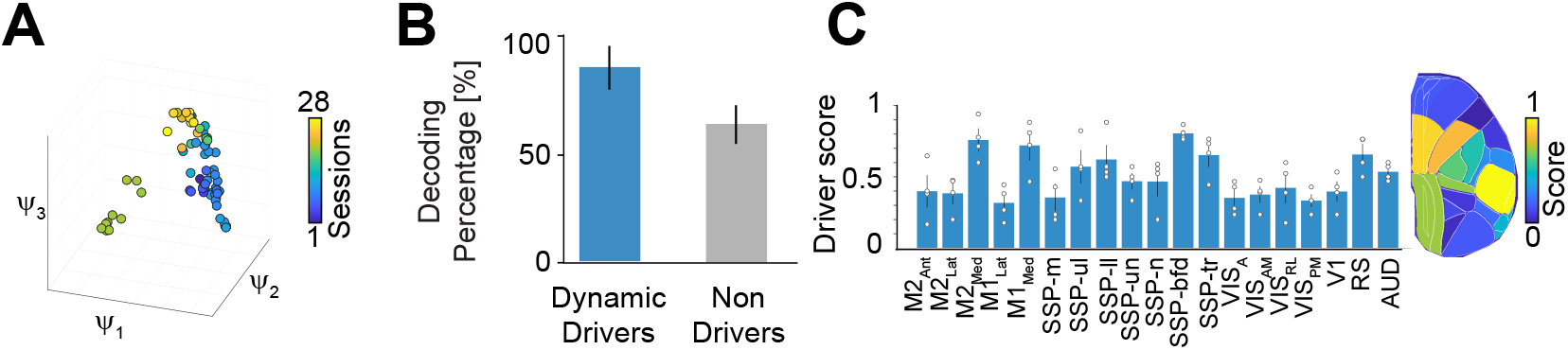
Cortical regions driving learning-related FC reorganization. (A) Cortex-wide FC trajectory during auditory-decision training visualized via diffusion-map embedding; color indicates session index (training progression). (B) Decoding power (percentage) calculated as the accuracy (*R*^2^) for predicting training stage (session index) from diffusion coordinates derived from the driving subnetwork, and non-driving subnetwork (mean *±* SEM, *n* = 4). (C) Driver scores highlight cortical regions that shape dynamics.

Finally, we aggregated driver scores across animals to obtain a region-level summary of contributions to learning-related FC dynamics. Driver scores consistently highlighted retrosplenial cortex, medial frontal areas, and specific somatosensory regions (Fig. 5C), suggesting that learning-related reconfiguration is dominated by a reproducible subset of cortical areas. Together, these results indicate that RONI isolates a stable set of regions whose time-varying interactions drive cortex-wide FC reorganization during learning.

### C. Dendritic dynamic drivers preferentially support emergence of task-relevant motor encoding

To probe FC dynamics at finer spatial scales, we analyzed dendritic calcium imaging from primary motor cortex (M1) during acquisition of a lever-pull task (8 mice; 15 training days; Fig. 6A). The same field of view was tracked across days, and ROIs were annotated along layer-5 apical tuft dendritic branches (about 12 cells per animal, see Appendix V H for more details). Although dendritic ROIs could be assigned to parent neurons based on anatomical reconstruction, we intentionally did not use this information as prior knowledge. For each trial, we computed the full pairwise correlation matrix across ROIs, yielding a trial-by-trial SPSD FC sequence per animal.

**FIG. 6.**
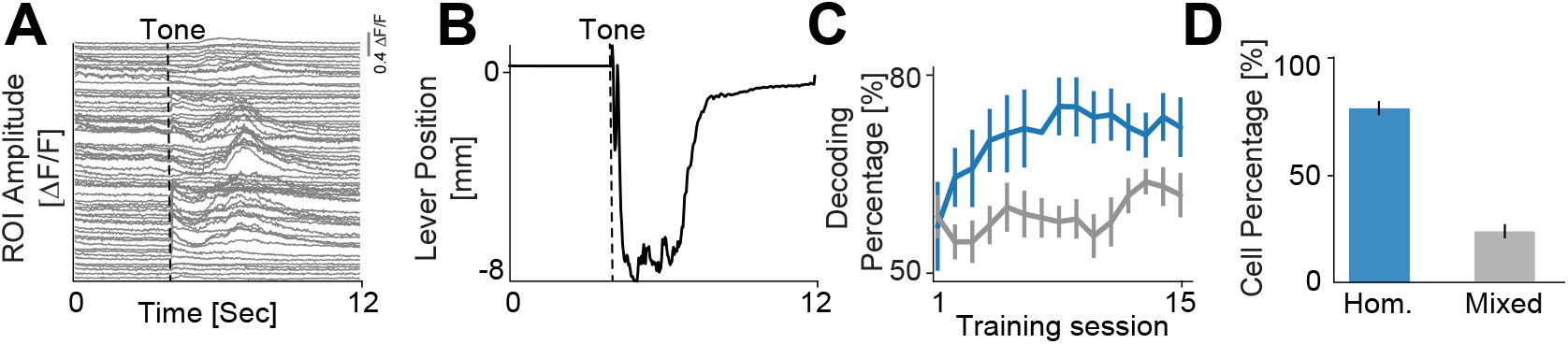
Dendritic subnetworks support task-relevant motor encoding. (A) Example dendritic activity from a single trial in an expert animal. A tone is given at 4 seconds, instructing the mouse to pull the lever. (B) Corresponding lever trajectory for the same trial as A in an expert animal. (C) encoding power [%] for predicting lever position from the activity of sub-populations (driver dendrites vs. non-drivers), normalized by the *R*^2^ achieved using the full neuronal population (mean *±* SEM, *n* = 8 animals). (D) Percentage of neurons classified as homogeneous, defined as having at least 80% of their dendrites either identified as dynamic drivers or as non-drivers (mean *±* SEM, *n* = 8 animals).

Applying RONI to these sequences identified a subset of dendritic ROIs as dynamic drivers in each animal. To test behavioral relevance, we trained linear models to predict lever position from dendritic activity and compared models trained with only driver ROIs with those trained with complementary non-driver ROIs (5-fold cross-validation). Across sessions and animals, driver-based models had greater decoding power than non-driver models (Fig. 6C), indicating that the subnetwork isolated by RONI preferentially supports task-relevant motor encoding during learning.

Moreover, driver labels were not uniformly distributed across dendrites. Grouping ROIs by parent neuron revealed a bimodal organization: most neurons were homogeneous, i.e., composed of driver or non-driver dendrites (Fig. 6). This organization suggests that the major sources of FC reconfiguration occur at the level of whole neurons, rather than being driven by isolated dendritic segments.

### D. RONI recovers contralateral sensorimotor structure in human EEG during motor imagery

Finally, we applied RONI to human EEG using a public motor-imagery dataset (BCI Competition IV, dataset IIa [47]) in which subjects imagined left-versus right-hand movement. Trials were band-pass filtered (8–30Hz), and correlation matrices were computed over a 3sec window starting 0.5sec before the go cue, yielding one SPSD matrix sequence per movement class.

We identified dynamic drivers separately for left- and right-hand imagery and aggregated driver scores across subjects. The resulting topography recapitulated the expected contralateral organization of motor control: electrodes over the left hemisphere contributed more strongly during right-hand imagery, whereas electrodes over the right hemisphere contributed more strongly during left-hand imagery (Fig. 7). These findings illustrate that RONI can extract an interpretable driver structure in human EEG that aligns with known neurophysiology.

**FIG. 7.**
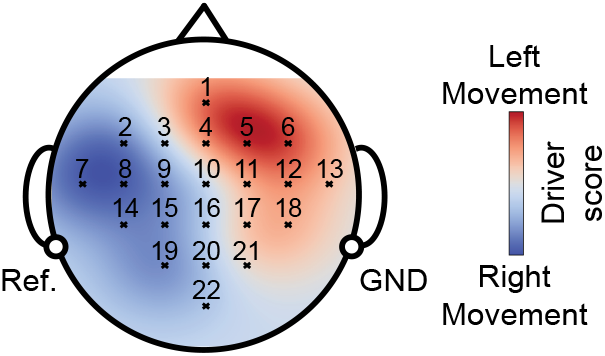
Contralateral sensorimotor organization in human EEG. Scalp maps of aggregated driver scores for left-(red) and right-hand (blue) imagery across subjects. Driver topographies exhibit contralateral dominance, matching known sensorimotor organization.

**FIG. 8.**
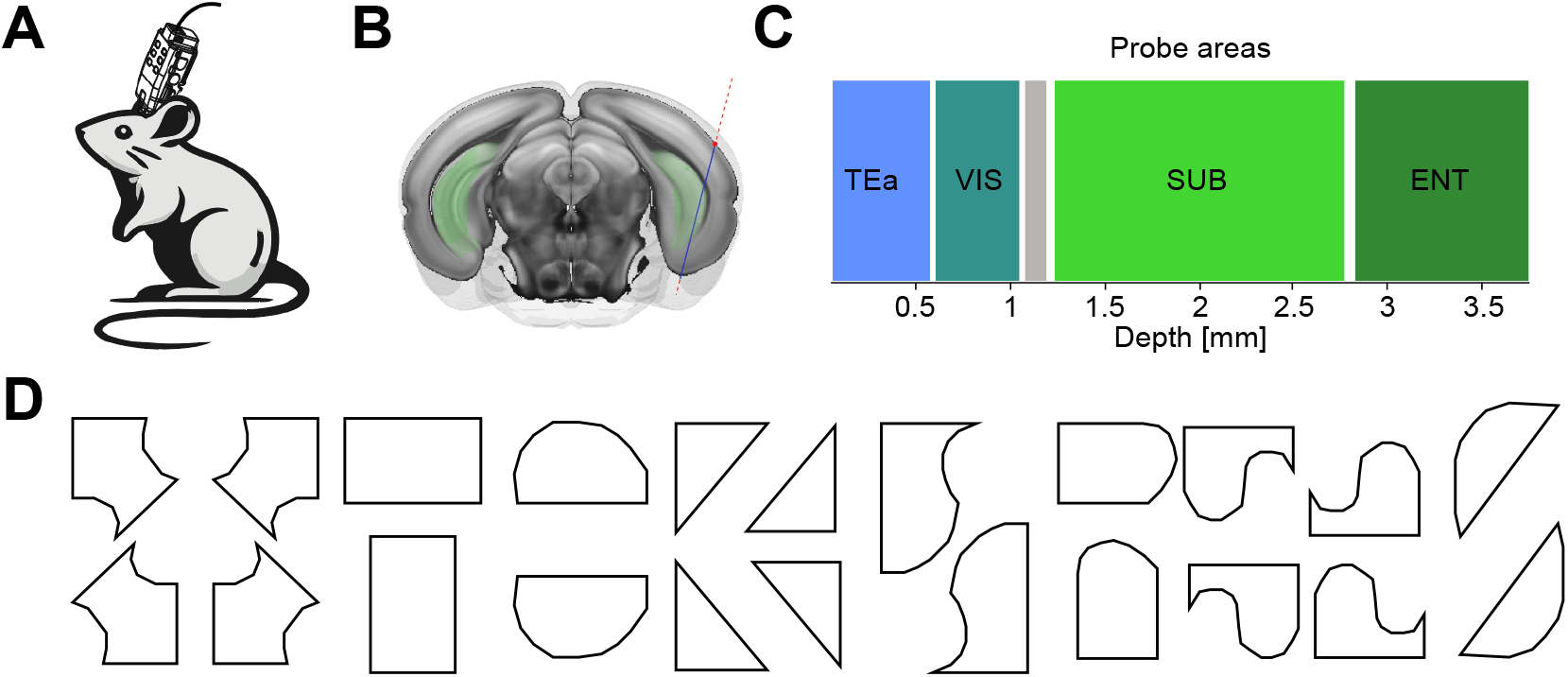
(A) Schematic of the mouse and Neuropixels probe used for electrophysiological recordings. (B) Coronal brain section showing the probe insertion site and recording trajectory. (C) Brain regions recorded during the experiment. (D) Outlines of all arena geometries used in the study.

## IV. CONCLUSION

We introduce RONI, a novel multi-resolution framework for analyzing temporal sequences of FC matrices that explicitly respects their underlying Riemannian geometry. By defining recursive low- and high-pass filtering operations directly on the manifold, RONI enables a multiresolution decomposition of structured correlation dynamics. Spectral analysis of the decomposed components enables the identification of dynamic drivers as sub-networks that shape temporal changes in the correlation structure. The successful application of RONI to very diverse experimental modalities and spatial scales demonstrates its robustness and general utility for studying neural population dynamics. Beyond neuroscience, RONI provides a general framework that could be extended to other domains for analyzing evolving structured data, including finance, climate science, and spatiotemporal graph learning. Future directions include addressing the method’s reliance on consistent components over time, a challenge in settings such as neuroscience, where tracking identical cells across sessions is often infeasible. Integrating our approach with deep learning may enable analysis of dynamic correlation structures with partial, noisy, or mismatched observations.

## V. APPENDIX

### A. Approximating the geodesic between two SPSD matrices

We summarize here the extension of the affine-invariant geometry from the SPD manifold to the SPSD cone following [45].

Let 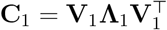 and 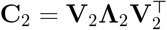 be two SPSD matrices of dimension *d* × *d*, where **V**_*k*_ ∈ ℝ^*d*×*r*^ contains orthonormal eigenvectors and **Λ**_*k*_ are diagonal matrices with positive eigenvalues.

First, we align the subspaces by computing the SVD 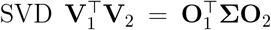, where **Σ** = diag(*σ*_1_, …, *σ*_*r*_), *r* = min (rank(**C**_1_), rank(**C**_2_)), and *θ*_*i*_ = arccos(*σ*_*i*_), and define **Θ** = diag(*θ*_1_, …, *θ*_*r*_), **U**_1_ = **V**_1_**O**_1_, and **U**_2_ = **V**_2_**O**_2_. We then interpolate the subspaces along the Grassmannian by defining

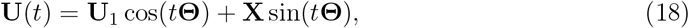

where 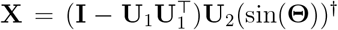 (sin(**Θ**))^*†*^ and (·)^*†*^ is the Moore–Penrose pseudoinverse. Next, we interpolate the SPD component by rotating the eigenvalue matrices 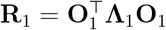 and 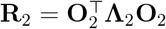, and forming the affine-invariant geodesic

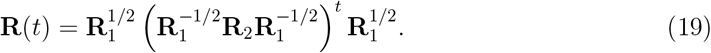

Finally, the approximate SPSD geodesic is reconstructed as

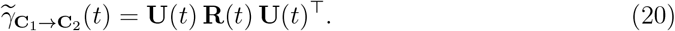

From this we can approximate the squared geodesic distance as:

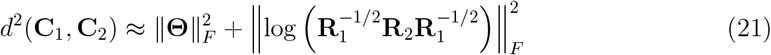

where the first term is the squared Grassmannian distance, and the second is the squared affine-invariant distance on the SPD manifold.

### B. Proofs for spectral analysis

The proof is given for SPD matrices. The result extends to SPSD matrices through the geodesic continuation, defined in Appendix V A.

#### Theorem 2.

Let *ψ* be an eigenvector of both **C**_1_ and **C**_2_ with eigenvalues *λ*_1_ and *λ*_2_, respectively: **C**_1_*ψ* = *λ*_1_*ψ*, **C**_2_*ψ* = *λ*_2_*ψ*. Then *ψ* is also an eigenvector of the difference operator **C**_1_%**C**_2_, with eigenvalue: 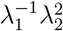.

*Proof*.

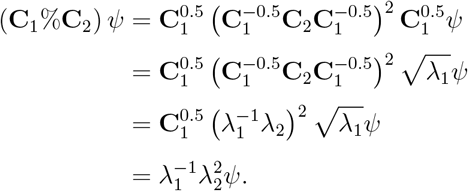

### C. Stability under weak perturbations

In practice, shared structure across matrices may only be approximate. The following analysis demonstrates that our operators are stable under small perturbations in shared eigenvectors, indicating that they behave well in the presence of noise. The *ε*-pseudospectrum of **C** is denoted Λ_*ε*_(**C**).

#### Definition 1

*Let* **C** ∈ ℝ^*d×d*^, *the following definitions of the* Λ_*ε*_(**C**) *are equivalent for a small ϵ* > 0 *[48]:*

1. *The set of λ* ∈ ℝ *such that* ∥(*λ***I** − **C**)^−1^∥ ≥ *ϵ*^−1^
2. *The set of λ* ∈ ℝ *such that λ* ∈ Λ(**C** + **E**) *for some* **E** ∈ ℝ^*d×d*^ *with* ∥**E**∥ ≤ *ϵ*
3. *The set of λ* ∈ ℝ *such that* ∥(*λ***I** − **C**)**v**∥ ≤ *ϵ for some* **v** ∈ ℝ^*d*^ *with* ∥**v**∥ = 1

The following two theorems extend our spectral results to the case of approximately common components, showing that pseudo-eigenstructure is preserved under our operators. We denote *S* = **C**_1_#**C**_2_ and *Q* = **C**_1_%**C**_2_.

#### Theorem 3

*[42] Suppose there exists an eigen-pair* (*λ*_*k*_, *ψ*_*k*_) *of* **C**_*k*_ *for k* = 1, 2 *so that ψ*_1_ = *ψ*_2_ + *ψ*_*ϵ*_, *where* 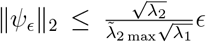 *for a small ϵ* > 0, *where* 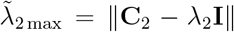 *Then we have:* 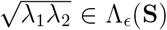. *Specifically, we have:* 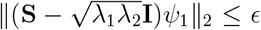, *and ψ*_1_ *is a corresponding ϵ–pseudo-eigenvector of* **S**.

#### Theorem 4

*Suppose there exists an eigenpair λ*_*k*_ *and ψ*_*k*_ *of* **C**_*k*_ *for k* = 1, 2 *so that ψ*_1_ = *ψ*_2_+ *ψ*_*ε*_, *where* 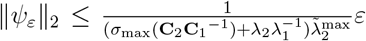, *where σ*_max_ *denotes the maximum eigenvalue, for a small ε* > 0, *and where* 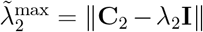. *Then we have* 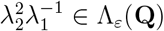. *Specifically, we have* 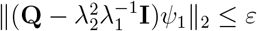, *and ψ*_1_ *is a corresponding ε-pseudo-eigenvector of* **Q**.

*Proof*. Using eqn. (23) from Appendix V E, we have:

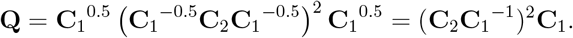

Since *ψ*_1_ is an eigenvector of **C**_1_ with eigenvalue *λ*_1_, we obtain:

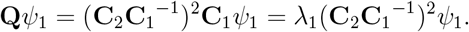

Thus, it suffices to show that *ψ*_1_ is an *ε*-pseudo-eigenvector of (**C**_2_**C**_1_^−1^)^2^. By direct expansion, we have:

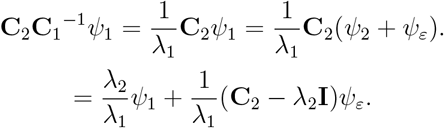

This implies:

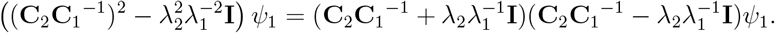

Multiplying both sides by *λ*_1_, we get:

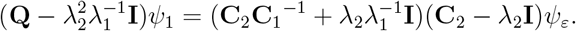

Taking the *ℓ*_2_ norm, we obtain:

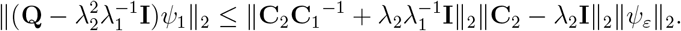

Applying the given bound on ∥*ψ*_*ε*_∥_2_, we conclude:

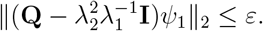

Thus, 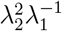 is an *ε* pseudo-eigenvalue of **Q**, where *ψ*_1_ is a corresponding *ε* pseudo-eigenvector. These results demonstrate that our operators are robust to perturbations in shared structure—a desirable property when working with noisy real-world data, such as neural recordings.

### D. Proof mutually inverse

#### Proposition 2.

Let **C**_1_ and **C**_2_ be SPD matrices, the operators defined in (4)(5) are mutually inverse, meaning:

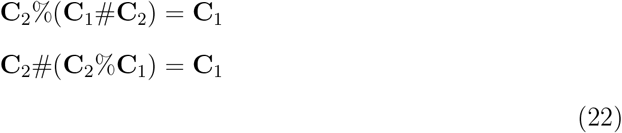

The same relations hold for SPSD matrices by continuity of the extended geodesic (Appendix V A).

*Proof*. We will first prove that **C**_2_%(**C**_1_#**C**_2_) = **C**_1_:

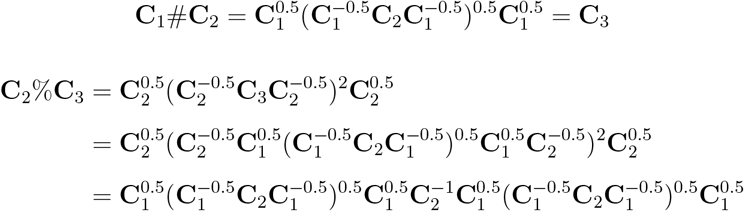

We now verify that this expression simplifies to **C**_1_:

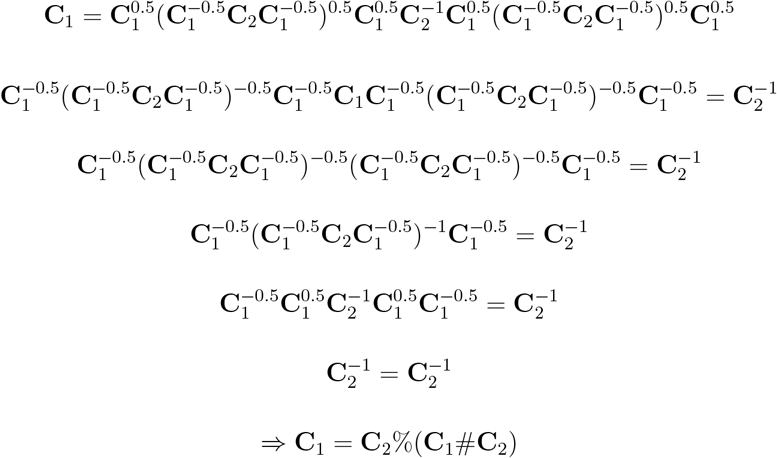

Now, we prove that **C**_2_#(**C**_2_%**C**_1_) = **C**_1_:

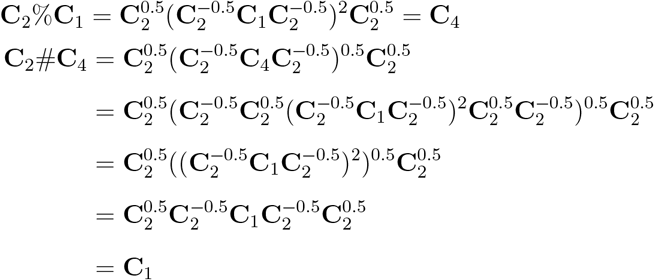

### E. Equivalent form

#### Proposition 3.

Equivalent form of the filters. We have

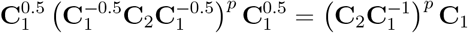

*Proof*. For some *p* ≥ 0, consider the expression:

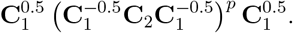

Define the matrices:

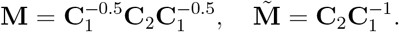

Since **C**_1_ and **C**_2_ are positive definite, so is **M**, which implies that **M** and 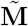 are similar via:

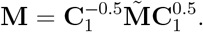

Let Λ^*M*^ and **V**^*M*^ be the eigenvalue and eigenvector matrices of **M**, respectively. Then the eigenvalue matrix of 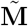 satisfies:

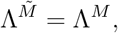

with right and left eigenvectors given by:

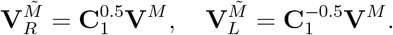

Thus, we obtain the matrix powers:

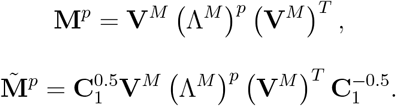

From this, we conclude:

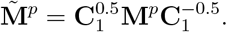

Therefore, we derive:

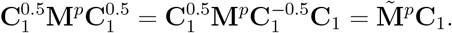

Substituting back for **M** and 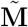, we obtain:

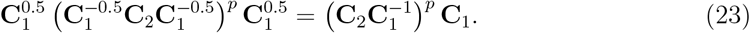

### F. The hippocampal network - navigation dataset

Neuropixels probes recorded neural activity from the subiculum, medial entorhinal cortex (MEC), and temporal association cortex (TEA) in freely moving mice. A motorized XY-stage tethering system enabled unrestricted, guided exploration for 20 min in a 60 *×* 60 *cm*^2^ arena. We considered 38 distinct geometries, four mice, and approximately 150 neurons per mouse. For each experiment, neurons were filtered by minimum mean rate ≥ 0.1 spikes per second, inclusion of Kilosort-classified “good” units [49], and exclusion of Bombcell-classified “noise” units [50].

### G. Cortical network during learning

Comprehensive details of the widefield imaging are provided in [46] and [51]. Widefield imaging was done with an inverted tandem-lens macroscope and a sCMOS camera (Edge 5.5, PCO) running at 30 frames per second and a field of view of 12.5 *×* 10.5 mm2 (Fig. 9A). Imaging resolution was 640 *×* 540 pixels after 4*×* spatial binning, resulting in a spatial resolution of about 20 *µ*m per pixel. Mice were Ai93D;Emx-Cre;LSL-tTA animals, expressing the calcium indicator GCaMP6f in all excitatory cortical neurons. All imaging data was then rigidly aligned to the Allen Mouse Brain CCF, using anatomical landmarks, to allow for inferring the activity of different cortical areas over the course of task learning (Fig. 9B). Accurate atlas alignment was further confirmed using retinotopic mapping, showing high agreement of functionally inferred locations of visual areas.

**FIG. 9.**
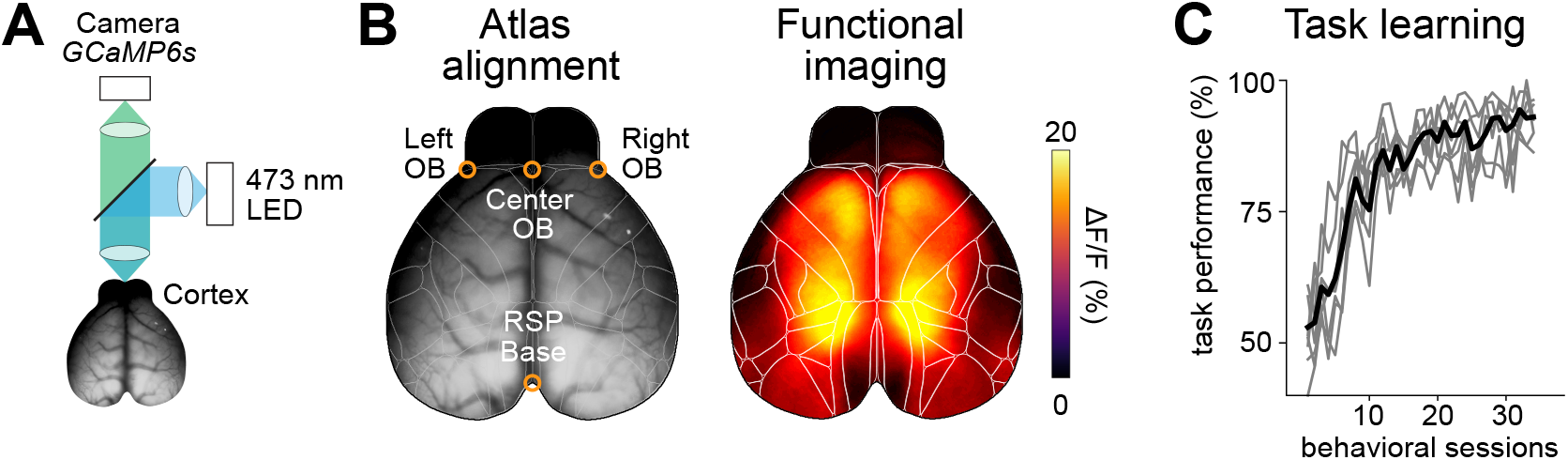
(A) Schematic of the widefield imaging setup for cortex-wide functional imaging. Imaging was performed through the intact, cleared skull. (B)Imaging data was aligned to the Allen CCF using 4 anatomical landmarks. Cortical activity, measured as relative changes in measured fluorescence, could then be related to the activity of cortical areas (white lines). (C) Learning curves of mice, trained in the auditory discrimination task. Animals achieved expert task performance within 10-20 sessions of behavioral training.

**FIG. 10.**
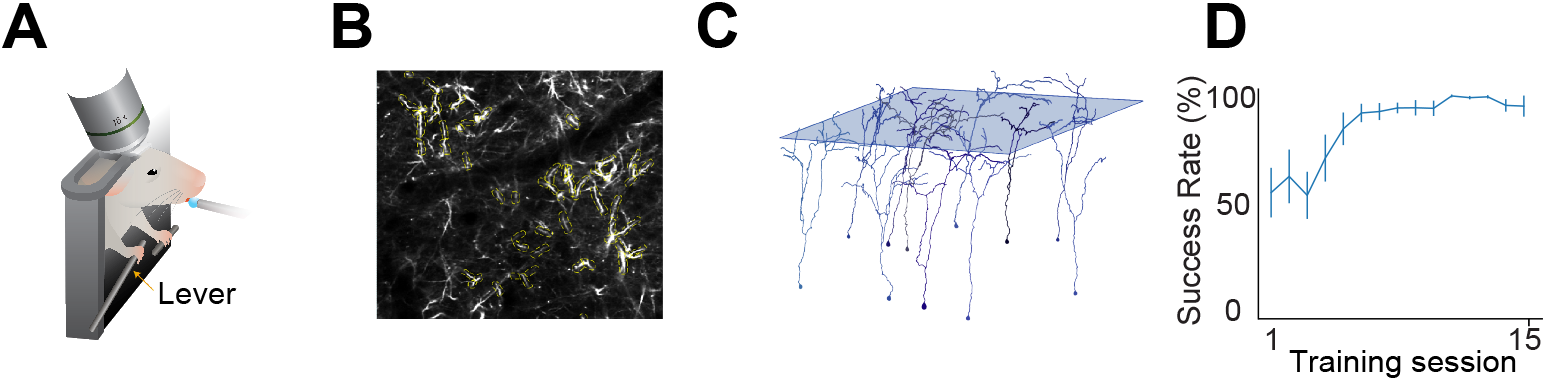
(A) Experimental setup: a head-fixed mouse performs a lever-pull task during two-photon calcium imaging. Each trial lasts 12 seconds, where a go-cue is given after 4 seconds. Lever position crossing the 5mm threshold is rewarded with a sweetened drop of water delivered by a water spout. (B) Example field of view showing manually labeled dendritic ROIs (yellow). (C) Morphological reconstruction and z-plane imaging. (D) Lever-pull success rate (mean±SEM) of 8 mice across training days, showing an increase in performance with training.

During widefield imaging, head-fixed mice were trained in an auditory discrimination task, where two sequences of Poisson-distributed click sounds were presented to either the left or right side of the animal for 1 to 1.5 seconds. After the stimulus and a subsequent 0.5-long delay period, mice had to perform a licking response on the side where more clicks were presented to obtain a water reward. The auditory discrimination task, therefore, required the accumulation of the auditory information over time and working memory to obtain high task performance (Fig. 9C). Animals were trained between 25 and 40 sessions, and we analyzed changes in cortical correlation structure over the course of task learning. Results here include cortex-wide population activity recorded from all training sessions of four animals.

### H. Dendritic networks in the primary motor cortex (M1) during motor learning

In this longitudinal experiment, eight mice were trained to perform a motor task during 15 sessions. Each trial lasted 12 seconds, during which an auditory go cue was given four seconds into the trial, prompting the animals to pull the lever. Pulls that surpassed a preset displacement threshold were rewarded by a drop of sweetened water delivered to the animal’s mouth through a water spout. Lever position was sampled at 150 Hz, yielding millisecond-resolution kinematic traces. Throughout every session, simultaneous two-photon calcium imaging (30 Hz) was conducted to monitor the apical tuft dendrites of 12 (on average) layer-5 Pyramidal Tract (L5 PT) neurons in M1 forelimb region per animal. For stable tracking of dendrites, a sparse labeling technique of L5 PT neurons was applied. In addition, the pons was injected at three depths with a viral mixture comprising (i) a diluted retrograde AAV carrying Cre recombinase, (ii) a concentrated retro AAV-CAG-FLEX-jGCaMP7s, and (iii) a concentrated retro AAV-FLEX-tdTomato. A chronic cranial window implanted over forelimb M1 enabled repeated imaging of the same field of view (FOV) across days. After each session, regions of interest (ROIs) were manually annotated, preserving the identity of individual dendritic segments across all recordings.

## Notes

### Competing Interest Statement

The authors have declared no competing interest.

### Summary of Updates

Added new results on electrophisiology data from the hippocampus.

### Funding

- Israel Science Foundation

